# Interpretable deep generative models for genomics

**DOI:** 10.1101/2021.09.15.460498

**Authors:** Yongin Choi, Ruoxin Li, Gerald Quon

**Affiliations:** Biomedical Engineering Graduate Group, University of California, Davis, Davis, CA; Genome Center, University of California, Davis, Davis, CA; Graduate Group in Biostatistics, University of California, Davis, Davis, CA; Department of Molecular and Cellular Biology, University of California, Davis, Davis, CA

## Abstract

Deep neural networks implementing generative models for dimensionality reduction have been extensively used for the visualization and analysis of genomic data. One of their key limitations is lack of interpretability: it is challenging to quantitatively identify which input features are used to construct the embedding dimensions, thus preventing insight into why cells are organized in a particular data visualization, for example. Here we present a scalable, interpretable variational autoencoder (siVAE) that is interpretable by design: it learns feature embeddings that guide the interpretation of the cell embeddings in a manner analogous to factor loadings of factor analysis. siVAE is as powerful and nearly as fast to train as the standard VAE but achieves full interpretability of the embedding dimensions. Using siVAE, we exploit a number of connections between dimensionality reduction and gene network inference to identify gene neighborhoods and gene hubs, without the explicit need for gene network inference. We observe a systematic difference in the gene neighborhoods identified by dimensionality reduction methods and gene network inference algorithms in general, suggesting they provide complementary information about the underlying structure of the gene co-expression network. Finally, we apply siVAE to implicitly learn gene networks for individual iPSC lines and uncover a correlation between neuronal differentiation efficiency and loss of co-expression of several mitochondrial complexes, including NADH dehydrogenase, cytochrome C oxidase, and cytochrome b.

## INTRODUCTION

Single cell genomic assays such as scRNA-seq and scATAC-seq measure the activity level of tens to hundreds of thousands of genomic features (genes or genomic regions), yielding high dimensional measurements of cells. Features tend to be inter-correlated: gene members of the same pathway, complex or module exhibit correlated expression patterns across cells^1^, and proximal genomic regions covering the same regulatory elements or expressed genes are correlated in their accessibility patterns^2^. Key analysis tasks such as visualization^3^, clustering^4^, trajectory inference^5,6^, and rare cell type identification^7,8^ typically do not directly compute on the original features. Instead, they first perform dimensionality reduction (DR) to project cells from their high dimensional feature space to a lower dimensional “cell embedding space” consisting of a smaller set of embedding dimensions. Individual embedding dimensions capture distinct groups of correlated input features, and are often also correlated with biological factors such as casecontrol status^9^, gender^10^, and others^11^. Downstream tasks are then carried out on these embedding dimensions.

Given the central role of embedding dimensions, it is useful to be able to characterize and interpret which of the original input features contributed to the construction of each embedding dimension. For example, in a visualization of a 2D cell embedding space, interpretation of the embedding dimensions would identify genes that explain variation in the transcriptome along different axes (**Fig. 1**). Linear DR frameworks such as PCA achieves interpretation through estimation of the contribution of individual features towards each embedding dimension. However, linear DR frameworks are considered less powerful because they can often be viewed as a restricted implementation of a non-linear framework^12^. In contrast, non-linear methods such as UMAP, t-SNE and variational autoencoders (VAEs) produce better visualizations in which cells of the same type cluster together more closely (**Supplementary Fig. S1**), but they are not interpretable and require more ad hoc, downstream analysis to gain intuition about the arrangement of cells in the visualization. Beyond visualization, interpretability is an important property for other tasks, such as the detection of genes and pathways driving variation in expression within or across cell types^13^ and identification of genes associated with cellular trajectories directly from visualization^14^.

**Figure 1:**
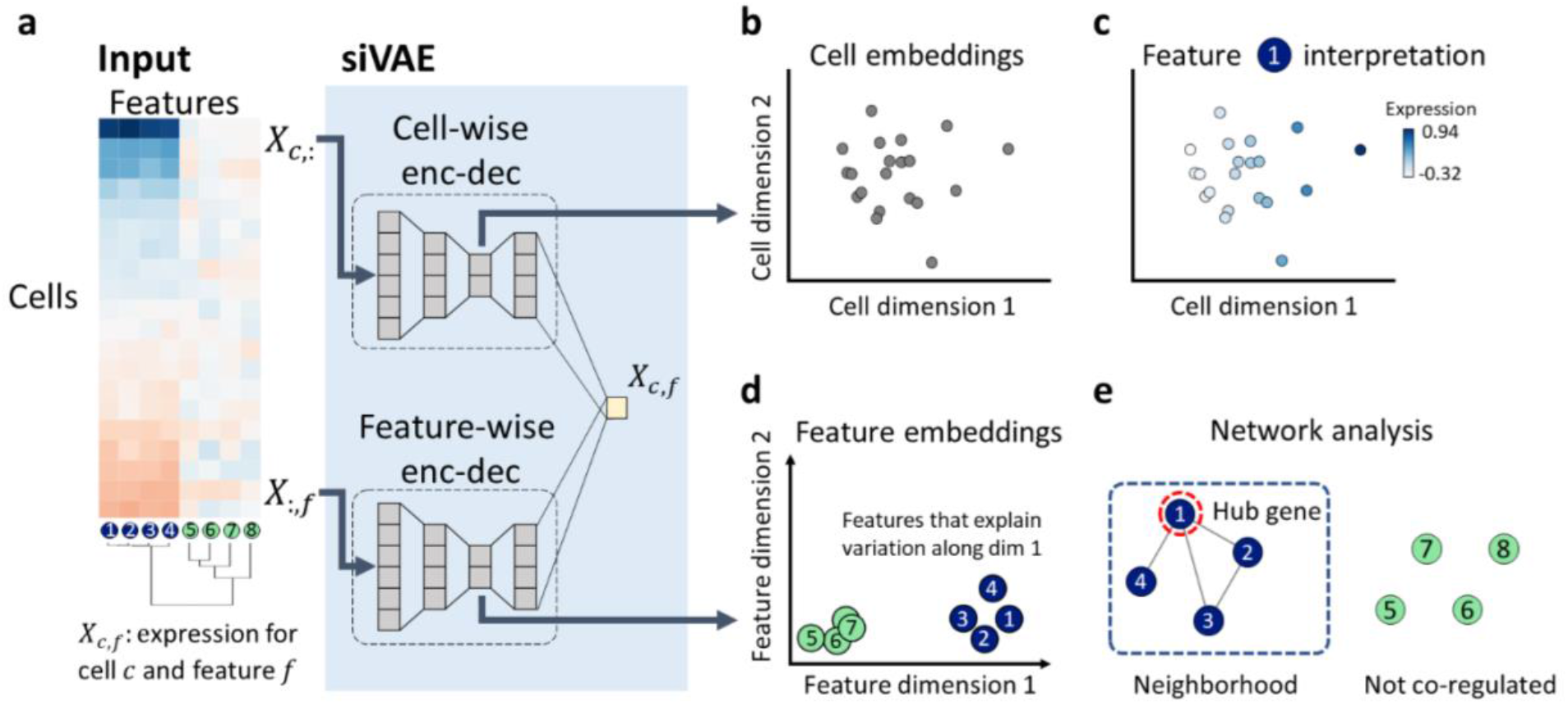
The siVAE model for inferring interpretable representations of single cell genomic data. (**a**) The input to siVAE is a cell by feature matrix; shown here is a synthetic gene expression matrix of eight genes, four of which are tightly regulated (genes 1,2,3, 4), and the other four of which vary independently (5,6,7, 8). siVAE is a neural network consisting of a pair of encoder-decoders, that jointly learn a cell embedding space and feature embedding space. The “cell-wise encoder-decoder” acts like a traditional VAE, where the input to the encoder is a single cell c’s measurement across all input features (*X*_*c*,:_). The cell-wise encoder uses the input cell measurements to compute an approximate posterior distribution over the location of the cell in the cell embedding space. The “feature-wise encoder-decoder” takes as input measurements for a single feature *f* across all input training cells (*X*_:,*j*_). The feature-wise encoder uses the input feature measurements to compute an approximate posterior distribution over the location of the feature in the feature embedding space. The decoders of the cell-wise and feature-wise encoderdecoders combine to output the expression level of feature *f* in cell *c* (*X_c,f_*). (**b**) Visualization of the cell and feature embedding space learned from the gene expression matrix in (a). Note the embeddings of genes 1,2, 3 and 4 all have large magnitudes along dimension 1 but not dimension 2, suggesting genes 1,2, 3 and 4 explain variation in the cell embedding space along dimension 1. Genes 5, 6, 7, and 8 sit at the origin of the feature embedding space, suggesting they do not co-vary with other features. (**c**) The expression patterns of gene 1 are overlaid on the cells in the cell embedding space. Gene 1 clearly increases in expression when inspecting cells from left to right, consistent with the feature embedding space that shows Gene 1 having large loadings on dimension 1. (**d**) In contrast, Gene 5’s expression does not have a clear pattern of variation with respect to position of the cell in cell embedding space, consistent with Gene 5’s location close to the origin in the feature embedding space. (**e**) A trained siVAE model can be used to identify hubs and gene neighbors in a gene co-expression network, without the need to explicitly infer a coexpression network.

Previous works have explored extensions of the non-linear VAE framework to achieve interpretability. Methods such as LDVAE^15^, scETM^16^, and VEGA^17^ achieve interpretability by imposing a linear relationship between the latent embedding layer and the output layer in the decoder, which in turn makes these approaches effectively linear dimensionality reduction methods. Other ad-hoc approaches (DeepT2Vec^18^ and deepAE^19^) combine unsupervised autoencoders with a supervised loss function that uses metadata such cell type information, which depends on having prior knowledge and restricts interpretation only with respect to those features.

Here we propose a scalable, interpretable variational autoencoder (siVAE) that combines the nonlinear DR framework of variational autoencoders with the interpretability of linear PCA. siVAE is a variant of VAEs that additionally infers a feature embedding space for the genomic features (genes or genomic regions) that is used to interpret the cell embedding space. Importantly, by using a non-linear network to combine the cell and feature embedding space, siVAE achieves interpretability without introducing linear restrictions, making it strictly more expressive than LDVAE, scETM and VEGA. Compared to other approaches for achieving interpretable, non-linear DR, siVAE is either faster, generates better low dimensional representations of cells, or more accurately interprets the non-linear DR without introducing linear restrictions or dependence on prior knowledge.

## RESULTS

siVAE is a deep neural network consisting of two pairs of encoder-decoder structures, one for cells and the other for features (**Fig. 1a**). The cell-wise encoder-decoder learns to compress percell measurements *X_c_*,. (where *X* is a matrix of dimension *C* × *F*, where *C* indexes cells and *F* indexes features) into a low dimensional embedding *(z_c_*) of length *K* for visualization and analysis, similar to traditional VAEs implemented in single cell genomic applications and others^20–22^. We call the *C* × *K* matrix of embeddings *Z* the siVAE score matrix, where the scores of cell *c* (*Z*_*c*,:_) represent its position in the cell embedding space.

To facilitate interpretation of the cell state space, siVAE additionally implements a separate feature-wise encoder-decoder network (**Fig. 1a**) that learns to compress per-genomic features across the cells (*X*_:,*f*_) into a low dimensional embedding *(v_f_*) of length *K,* analogous to the cellwise encoder-decoder. We call the *F* × *K* matrix of feature embeddings *V* the siVAE loading matrix, where the loadings of feature *f*(*V*_*f*,:_) represent its position in the feature embedding space. The cell- and feature-wise decoders together are used to generate the observed measurement *X_C,f_*.

The strategy siVAE uses to achieve interpretation is best understood by briefly reviewing why probabilistic PCA (PPCA) and factor analysis are interpretable^15,23^. The underlying generative model behind PPCA can be thought of as similar to a VAE with a linear decoder, and the output of PPCA includes both a factor loading matrix *V* and score matrix *Z*. In probabilistic PCA, the predicted expression of feature *f* in cell *c* (*X_c,f_*) is assumed to be 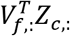, the dot product of the loadings for feature *f* and the scores of cell *c*. PPCA is therefore interpretable, because the larger the contribution of a feature *f* to a particular dimension *k* (indicated by the magnitude of *V_f,k_*), the more the measurement of feature *f* (*X_c,f_*) is influenced by a cell’s corresponding score in that dimension (*Z_c,k_*). Conversely, when the magnitude of *V_f,k_* is small (or even 0), then the cell’s corresponding score in that dimension (*Z_c,k_*) does not influence *X_c,f_*, the measurement of feature *f* in cell *c.* In this regard, we say that the PPCA model enforces correspondence between *Z_c,k_* and *V_f,k_*, the cell and feature embedding at dimension *k*.

siVAE achieves interpretability of the siVAE scores *Z_c,k_* by adding a small interpretability regularization term to its objective function (see Methods). More specifically, this regularization term penalizes deviation between the observed measurement *X_c,f_*, and the dot product of the corresponding siVAE scores and loadings 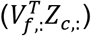. This small regularization term helps enforce some soft correspondence between dimension *k* of the cell scores, and dimension *k* of the feature loadings.

Our framework for making VAEs interpretable is generalizable to other VAE-based frameworks. Given that VAEs have been applied to a wide range of genomics data modalities (epigenomics^24–26^ and miRNA^27^) and analysis (visualization^20–28^, trajectory inference^29^, data imputation^30^, and perturbation response prediction^31–33^), our work can therefore enable interpretability in a wide range of downstream applications of VAEs.

## RESULTS - siVAE accurately generates low dimensional embeddings of cells

We first evaluated siVAE in the context of cell embedding space inference, where the goal is to generate low dimensional representations of cells in which cells of the same cell type cluster together. We benchmarked siVAE against other interpretable and non-interpretable dimensionality reduction approaches using a fetal liver cell atlas^34^ consisting of 177,376 cells covering 41 cell types. We measured the accuracy of each approach in a 5-fold stratified cross validation framework by first using the training folds to learn a cell embedding space, followed by training of a k-NN (*k* = 80) classifier using the known cell type labels and cell coordinates within the embedding space. We then classified the held-out cells. We associate higher k-NN accuracy with a more accurate cell embedding space in which cells of the same type cluster together.

We compared siVAE against a classic VAE as well as LDVAE^15^, where all three VAE frameworks used cell-wise encoder-decoders of the same size, and the VAE and siVAE use the same activation functions. Overall, we found siVAE’s cell embedding space to be comparable in accuracy to classic VAEs, suggesting that the introduction of the siVAE feature-wise encoderdecoder does not affect siVAE performance in terms of its cell embedding space. 2D visualization of siVAE’s cell embedding space reveals strikingly similarity to the cell embedding space of the classic VAE in that cells of the same type cluster together (**Fig. 2a**). Furthermore, siVAE is competitive in classification accuracy with a classic VAE on the fetal liver cell atlas (**Fig. 2b**). siVAE therefore is competitive with VAEs in terms of generating cell embedding spaces, but has the additional benefit of interpretability, which we will explore below. In comparison, the LDVAE approach, which is interpretable like siVAE but performs linear DR, yields significantly lower classification accuracy (**Fig. 2b**) and generates visualizations in which different cell types mix together more prominently (**Fig. 2a**). LDVAE therefore gains interpretability at a cost to the accuracy of the cell embedding space.

**Figure 2:**
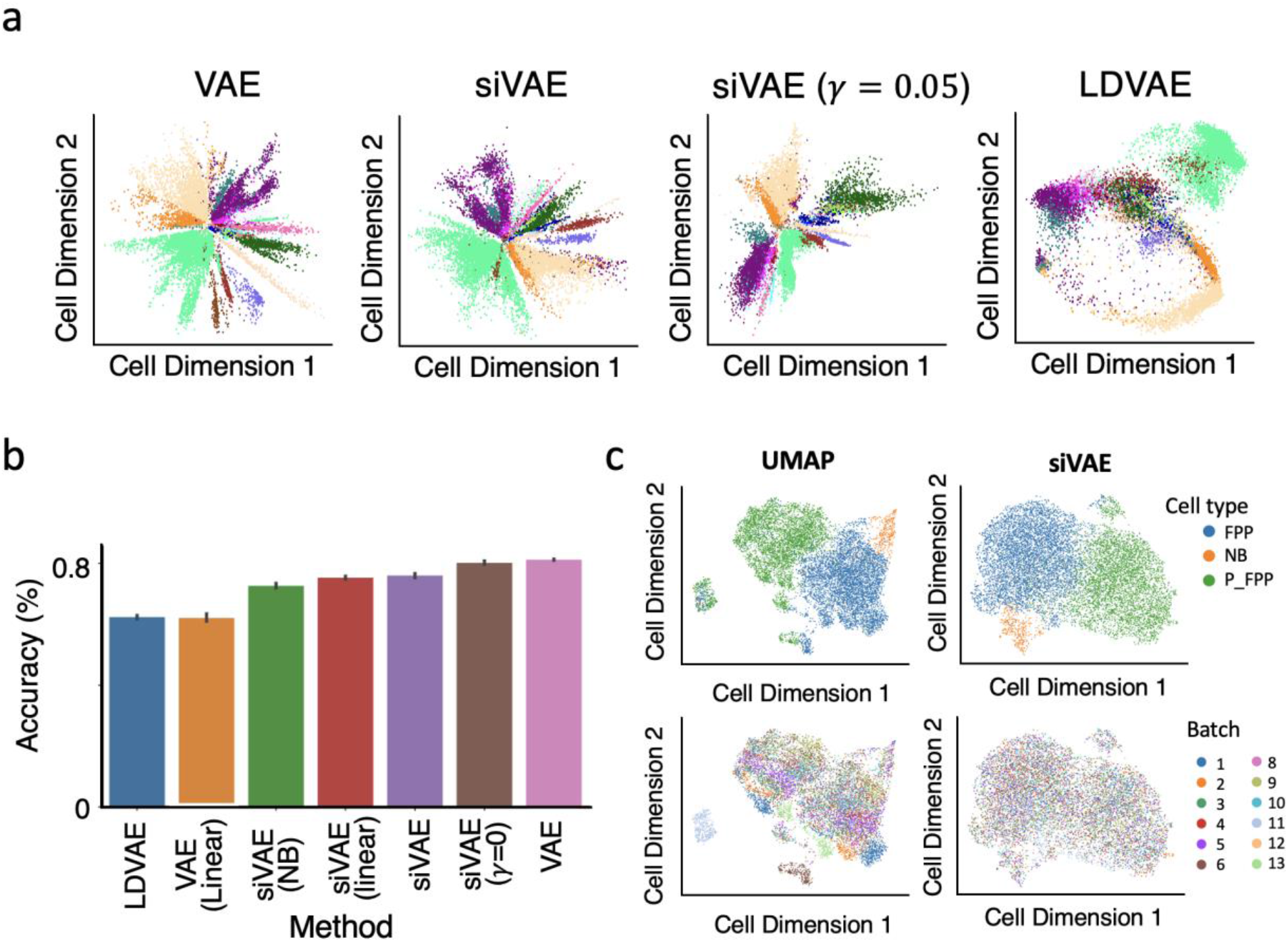
Accuracy evaluation of siVAE’s cell embedding spaces. (**a**) 2D visualization of the inferred cell embedding spaces of a classic VAE, siVAE, a variant of siVAE in which the interpretability regularization term is removed (*γ* = 0) and LDVAE. Each point represents a cell and is colored according to cell type. (**b)** Barplot indicating the accuracy of a *k*-NN (*k* = 80) classifier predicting the cell type labels of single cells based on their inferred position in the cell embedding space inferred by siVAE and other methods trained on the Fetal Liver Atlas dataset. (**c**) 2D visualization of the inferred cell embeddfing space using UMAP and siVAE with batch correction. Top row shows annotation based on cell type, and the bottom row shows annotation based on batch.

We next constructed a set of model variants of siVAE in order to identify which aspects of siVAE lead to its superior performance over LDVAE (**Table 1**). LDVAE is in principle similar to the classic VAE, with two key differences. First, the LDVAE decoder is restricted to use only linear activation functions in order to achieve interpretability; thus, LDVAE performs linear dimensionality reduction. Second, the LDVAE loss function uses a negative binomial or zero-inflated negative binomial distribution over the input features (genes), instead of the Gaussian distribution used in a classic VAE. In principle, the NB or ZINB observation model is a better fit for single cell transcriptomic data compared to a Gaussian distribution normally used on log transformed data ^35,36^. We therefore constructed two variants of siVAE, siVAE-NB and siVAE-linear. siVAE-NB is identical to siVAE, except that it uses a negative binomial distribution for the observation layer while maintaining non-linear activation functions in its decoders to achieve non-linear DR. siVAE-linear is identical to siVAE, except that it restricts both the feature-wise and cell-wise decoder to use linear activation functions like LDVAE and does not implement the interpretability term. **Fig. 2b and Supplementary Fig. S2** shows that siVAE-NB performs worse than the corresponding model with the Gaussian distribution (siVAE), suggesting that using a NB output layer does not lead to a more accurate cell embedding space. siVAE-linear is more accurate than LDVAE (**Fig. 2b**), indicating that the feature-wise encoder-decoder of siVAE is overall beneficial to dimensionality reduction. However, siVAE-linear performs more poorly than siVAE, verifying the non-linear activation functions are beneficial to dimensionality reduction.

**Table 1.** List of model variations with corresponding features. Usage of the interpretability term only applies to siVAE and its variants. A linear decoder is composed of the same number of layers as the non-linear decoder unless specified as single, in which case the latent embedding layer is directly transformed to an output layer.

| Model | Interpretability Reg. | Decoder | Observation model |
| --- | --- | --- | --- |
| siVAE | Yes | Non-Linear | Gaussian |
| siVAE ($\gamma = 0$) | No | Non-linear | Gaussian |
| siVAE-linear | No | Linear | Gaussian |
| siVAE-NB | Yes | Non-linear | Negative Binomial |
| VAE | NA | Non-linear | Gaussian |
| scVI | NA | Non-linear | Negative Binomial |
| LDVAE | NA | Linear, single | Negative Binomial |
| VAE (linear) | NA | Linear, single | Negative Binomial |

We also hypothesized that the interpretability term used in siVAE’s loss function would degrade the quality of dimensionality reduction to an extent, as siVAE uses the regularization term to enforce correspondence between the individual dimensions of the feature and cell embedding spaces to achieve interpretability. We therefore constructed siVAE (γ=0), representing a siVAE model in which we turn off the regularization term by setting its weight *γ* to 0 and therefore disable interpretation, but keep the feature embedding space. From **Figure 2b**, we can see the small gap in classification performance between siVAE and VAE closes with siVAE (γ=0), showing that the intepretability of siVAE comes at a small cost in classification performance, though not nearly as large a cost as using linear dimensionality reduction, as evidenced by the poorer performance of siVAE-linear and LDVAE. We performed additional experiments to show that when varying *γ* from 0 to 100, where siVAE *(γ* =100) conceptually behaves similarly to siVAE-linear, the performance of siVAE smoothly interpolates between siVAE (*γ* =0) to siVAE-linear’s performance (**Supplementary Fig. S4**). This suggests that siVAE can be used to carefully balance interpretability with non-linear dimensionality reduction capability.

Finally, siVAE natively allows batch correction within the model similar to other models^16,20^. We tested our model on an iPSC neuronal differentiation (NeurDiff) dataset^37^ in which XX cell iPSC cell lines were sequenced using 10x Chromium before and after initiation of differentiation into neurons. We specifically focused on the samples from Day 11 before differentiation, as we observed a strong batch effect with respect to pool_id (**Fig. 2c**). When we trained siVAE and provided batch information during training, clustering by batch is eliminated while the clustering by cell type is still preserved (**Fig. 2c**).

## RESULTS - siVAE interprets cell embedding spaces faster and more accurately than existing feature attribution approaches

Having shown siVAE generates cell embedding spaces competitive with classic VAEs, we next verified that the interpretations of the embedding dimensions output by siVAE are accurate. Again, we define an interpretation of the cell embedding space as a matrix of feature loadings (or more generally, attributions) *V* of size *F × K,* where *F* is the number of features (e.g. genes), *K* is the number of cell dimensions, and the magnitude of *V_f,k_* indicates the strength of association between cell state dimension *k* and feature *f* in the original data space.

In contrast to methods such as siVAE and LDVAE that construct interpretable cell embedding spaces by design, there are two competing types of approaches to feature attribution in the literature that can help interpret cell embedding spaces post-inference. First, siVAE feature embeddings are analogous to general neural network feature attribution methods that quantify how each output node of a neural network depends on each input node (feature) of the network^38^, and include methods such as DeepLIFT^39^, saliency maps^40^, grad x input^39^, integrated gradients^41^, Shapley value^42^ and others^38,43–47^. One of the strengths of these approaches is they can be applied to any trained neural network in principle, making them highly generalizable. Second, methods such as Gene Relevance^48^ have been developed specifically to interpret cell latent spaces for any DR method including those not based on neural networks, and can be applied after cell embedding spaces are learned.

We first compared siVAE against Gene Relevance, using the neural network feature attribution methods as a gold standard as they have been extensively validated in other applications^49^. **Figure 3a** shows the mean pairwise correlation between the attributions of siVAE, Gene Relevance as well as three neural net feature attribution methods (saliency maps, grad x input, and DeepLIFT), where correlations have been averaged over each of the two feature dimensions that siVAE used to infer the cell embedding space for the fetal liver dataset. We see siVAE loadings are highly correlated with the neural net feature attribution methods (median Spearman *ρ* =0.73, P=1.1×10^-15^) with siVAE in striking agreement with DeepLIFT in particular (median Spearman correlation of 0.98, P=2.2 x10^-16^). In contrast, while Gene Relevance produced feature attributions that were consistent across their parameter selections (median Spearman correlation of 0.84, P=3.10 x10^-22^), they were poorly correlated with both neural net feature attribution methods (median Spearman correlation of 0.11, P=2.1×10^-6^) and siVAE (median Spearman correlation of 0.14, P=3.9 x10^-6^). These results suggest Gene Relevance is less accurate compared to siVAE at interpreting cell embedding spaces of VAE architectures. We found consistent results when comparing these methods on the MNIST imaging dataset (Supplementary Note 1).

**Figure 3:**
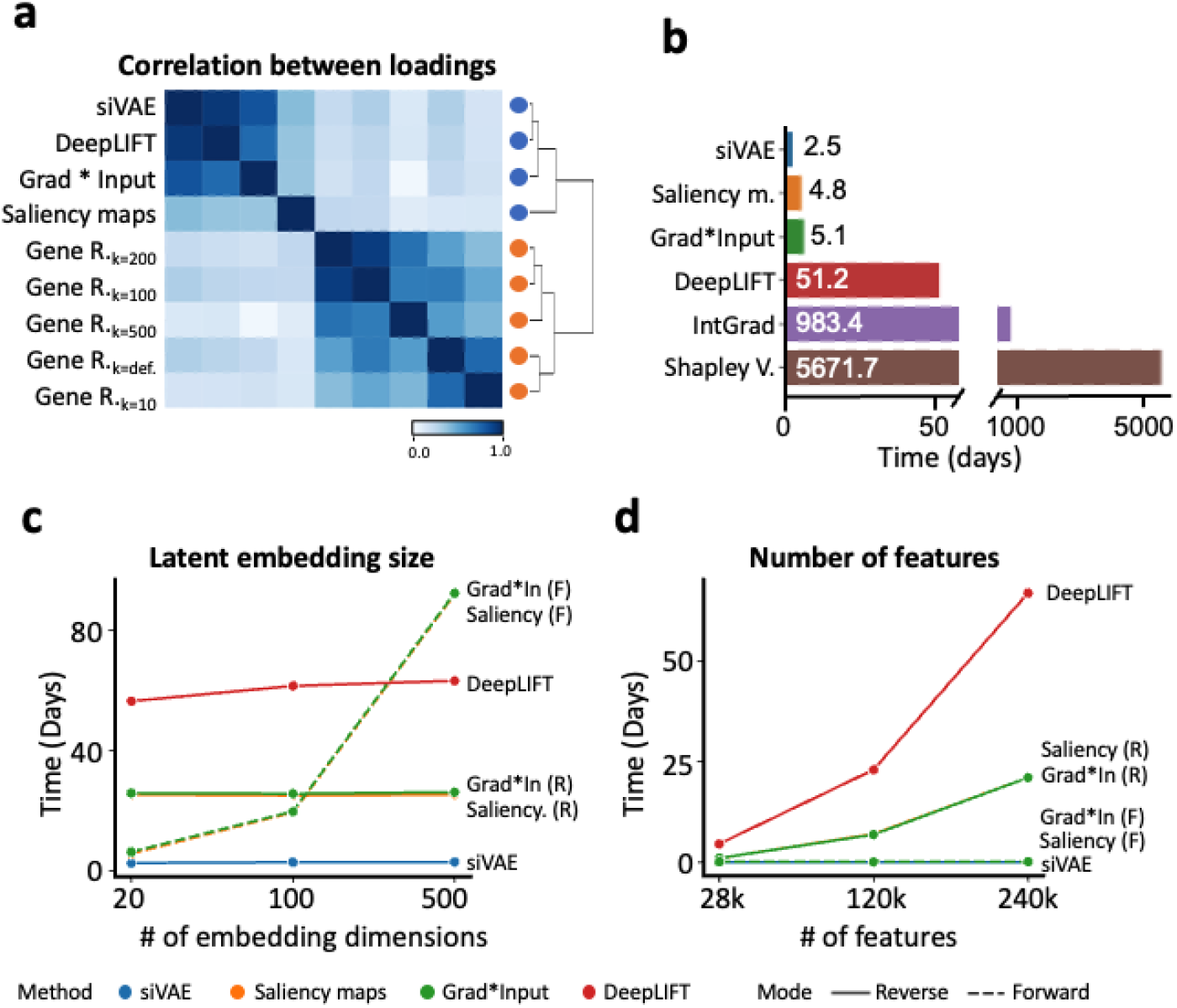
siVAE yields accurate and fast interpretations. (**a**) Heatmap indicating the mean pairwise correlation between the interpretations (loadings) of siVAE, Gene Relevance as well as three neural net feature attribution methods (saliency maps, grad*input, and DeepLIFT), where correlations have been averaged over each of the 2 embedding dimensions for the fetal liver atlas dataset. (**b**) Barplot indicating the time required to train siVAE versus training a classic VAE and applying feature attribution methods on the LargeBrainAtlas dataset. (**c**) Line plot indicating the time required to train siVAE and feature attribution methods on the LargeBrainAtlas dataset when the number of embedding dimensions for siVAE is varied, and the number of features is fixed at 28k. (**d**) Line plot indicating the time required to train siVAE and feature attribution methods on the scATAC-Seq dataset when the number of features is being varied, while the number of embedding dimensions is fixed at 20.

For the above results, we applied the neural net attribution methods to the decoder of siVAE to generate the ground truth feature attributions. Previous work has suggested to apply attribution methods to the encoder to improve execution speed^19,22^. Here we found that running attribution methods on the siVAE encoder produce substantially different interpretations that are in strong disagreement with interpretations of the decoder (**Supplementary Fig. S6**), suggesting interpretation of the encoder is not appropriate. These results make sense considering the primary role of the encoder is to compute an approximate posterior distribution of the latent embedding of each cell, as opposed to the decoder, which is responsible for directly mapping points from the latent space to gene expression space. Our results therefore suggest feature attributions should be applied to the decoder of VAEs instead of the encoder.

During our experiments on interpreting cell embedding spaces, it became evident that a number of neural network feature attribution approaches were computationally expensive to execute. Because these feature attribution methods perform calculations separately for either each embedding dimension or each output node of the network, their run time scales linearly with the number of embedding dimensions or features when run on VAE decoders^20^. The number of embedding dimensions is expected to be larger as the number of cells in the dataset grows, to accommodate more heterogeneity in the dataset; also, the number of features would be expected to be large for assays such as scATAC-seq that profile hundreds of thousands of genomic regions or more. We conjecture this problem of long execution time has not been previously reported in the literature because feature attribution methods are typically run on supervised neural networks to interpret class label predictions, and so the number of output nodes is traditionally very small, unlike generative models in genomics applications.

We therefore hypothesized that siVAE scales faster than the neural network attribution methods on larger single cell genomic datasets. To test this hypothesis, we assembled two datasets for execution time testing: the LargeBrainAtlas dataset published by 10x Genomics^50^ consisting of 1.3 million brain cells and 27,998 genes measured with scRNA-seq, and the BrainCortex dataset^51^ consisting of 8k cells and 244,544 genomic regions measured with SNARE-seq. We first compared the execution time of training siVAE on the full LargeBrainAtlas dataset, against the run time of training a VAE and individually running each of five neural network attribution methods (saliency maps, grad*input, DeepLIFT, integrated gradients and Shapley values) on the trained VAE. We found that siVAE achieved an execution time of 2.5 days, less than half of the fastest neural network attribution method (forward mode of saliency maps) (**Fig. 3d**).

To identify the most time-consuming aspects of feature attribution calculations for each method, we selected a subset of the LargeBrainAtlas dataset for varying the number of embedding dimensions from 20 to 512 and a subset of BrainCortex dataset for varying the number of features from 28k to 240k, to identify the speed bottlenecks. siVAE averaged 0.0073 days per embedding dimension (**Fig. 3e**) and 0.0027 days per 10k features (**Fig. 3f**), indicating siVAE execution time was robust to both the number of cells and features. On the other hand, we found the neural network attribution methods scale well when either the number of embedding dimensions or the number of input features is large, but not when they are both large. For example, DeepLIFT, Grad*Input (reverse-mode) and Saliency maps executed at 0.014, 0.0053, and 0.0012 days per embedding dimension respectively (**Fig. 3e**), but scaled poorly with respect to number of input features and executed at 2.9, 0.95, and 0.94 days per 10k features respectively (**Fig. 3f**). Switching Grad*Input and Saliency Maps to forward-mode led to fast execution times with respect to the number of input features (5.3×10^-4^ and 6.5 x10^-4^ days per 10k features respectively (**Fig. 3f**)) but led to poor scaling with respect to the number of embedding dimensions (0.18 and 0.17 days per embedding dimension, respectively (**Fig. 3e**)). Slower attribution methods such as Integrated Gradients and Shapley Value were excluded due to their infeasible execution times. In summary, the neural network attribution methods scale poorly either with the number of embedding dimensions or the number of input features depending on whether forward- or reverse-mode is used. This therefore makes their execution time slow relative to siVAE if both the number of features and embedding dimensions is large.

## RESULTS - Co-expressed genes cluster in the feature embedding space

Feature attributions, or factor loadings, of linear DR methods such as PCA have been exploited extensively in the literature to gain insight into the structure of gene co-expression networks (GCN)^15,23,52^; here we explore the extent to which the siVAE loading matrix can be leveraged to gain insight into GCN structure. GCNs are graphs in which nodes represent genes and edges represent co-expression of a pair of genes. A GCN captures co-variation in gene expression measurements between pairs (or more) of genes across a population of cells. GCNs are of interest because they can be used to identify (1) cell population-specific gene modules, representing groups of genes that are highly co-expressed and therefore are likely to function together in a cell type-specific manner, as well as (2) gene hubs, which are genes that are connected to an unusually large number of other genes, and typically represent key functional genes in the cell^53,54^. While GCN inference is valuable for interrogating gene regulatory patterns in a cell, GCN inference is a notoriously difficult and error-prone task^55–57^.

In our application of dimensionality reduction in which features are all centered and scaled uniformly, the goal of DR methods is to learn (linear or non-linear) patterns of co-expression amongst the input features, that allow accurate reconstruction of the input data from low dimensional representations. It is therefore natural to ask whether a trained siVAE model could yield insight into gene co-expression network structure of the training data, without the need for explicit gene network inference. More specifically, we view the siVAE loading matrix that siVAE infers as a non-linear analog of the PCA loading matrix. Indeed, one can view probabilistic PCA^58^ as a restricted form of a VAE in which all of the activation functions in the decoder are linear, no regularization is applied to the decoder weights, and the output distribution is an isotropic Gaussian.

Previous work has shown that eigengenes (genes captured by factor loadings of PCA) represent network modules in the gene co-expression network^59,60^. We hypothesized that siVAE genes captured by feature loadings of siVAE may also represent network modules, and that coexpressed genes in the training data are also proximal in the siVAE feature embedding space. To explore how a group of co-expressed genes are organized in the feature embedding space, we constructed a synthetic gene regulatory network consisting of five communities of 50 tightly correlated genes each, as well as an additional group of 50 independent, isolated genes (**Fig. 4a**). Each community follows a hub-and-spoke model in which a hub gene is connected to every other gene in the community, and each gene in the community is in turn only connected to the hub. No edges connect genes from different communities. Based on this gene network, we sampled a single cell gene expression dataset consisting of 5,000 cells and 300 genes (see Methods). The sampled expression matrix was used to train siVAE to embed genes in its feature embedding space.

**Figure 4:**
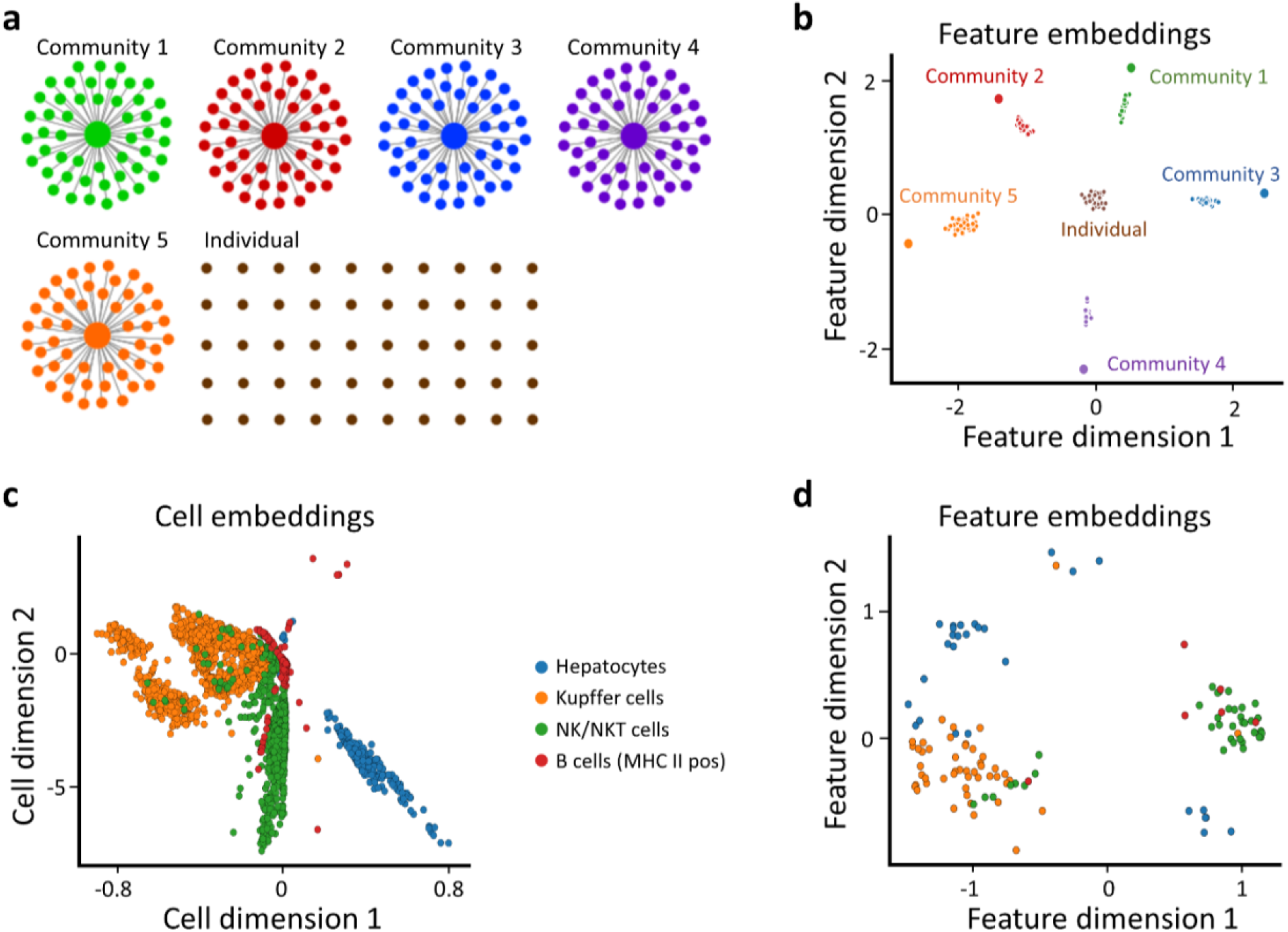
Co-expressed genes tend to co-localize in the siVAE feature embedding space. (**a**) The gene co-expression network used to simulate single cell expression data. The network consists of five tightly correlated groups of 50 genes each, along with 50 isolated genes. (**b**) Scatterplot of the feature embeddings produced from siVAE trained on the dataset simulated from the network in (a). Each point represents a gene, colored and labeled by the community it belongs to in (a). (**c**) Scatterplot of the cell embedding space inferred by siVAE trained on the fetal liver atlas dataset. Each point represents a cell and is colored based on its pre-defined cell type. (**d**) Scatterplot of feature embeddings inferred by siVAE trained on fetal liver atlas dataset. Each point represents a marker gene and is colored based on its prior known association to a cell type.

We found that genes belonging to the same community co-localized in the feature embedding space, but interestingly, the hub nodes are embedded in distinct locations their corresponding community (**Fig. 4b**). Our interpretation of this observation is that given the limited capacity of the cell embedding space, siVAE tends to keep information specifically about each hub because of their high degree centrality. On the other hand, non-hub genes within the same community colocalize in the feature embedding space because the limited capacity of the VAE forces non-hub genes to be predicted similarly, given the retained information about the hub. Interestingly, the 50 independent, isolated genes in the network were clustered tightly but near the origin in the feature embedding space, whereas genes that are part of a community are clustered but located farther away from the origin. This is likely because of two reasons. First, the KL divergence term of the feature-wise encoder-decoder of siVAE will tend to draw genes towards the origin. Second, because isolated genes by definition do not co-vary with other genes, information about their expression pattern will tend to be lost during compression, leading the VAE to tend to predict the average expression level of that gene in the decoder (which will be 0, because of data centering). This in turn encourages the feature embedding to be at the origin because the interpretability term encourages the linear product of the feature embedding with the feature loadings to predict the gene’s expression pattern, so if a feature is located at the origin in the feature embedding space, it will cause the predicted expression to be 0.

We also confirmed that co-expressed genes cluster in the feature embedding space using the fetal liver cell atlas data. Unlike the simulations above, for the fetal liver atlas there are no groundtruth gene regulatory networks available to use to identify truly co-expressed genes that are part of the same underlying gene communities. We therefore trained siVAE on the entire fetal liver atlas with 40 cell types, and considered marker genes of the same cell type^61^ to be a ground truth set of co-expressed genes. We selected only the four cell types with available marker genes for visualization (**Fig. 4c**). In the resulting feature embedding space learned by siVAE, we see that markers of the same cell type tend to cluster in feature embedding space as expected (**Fig. 4d**). Our results overall suggest that co-expressed genes tend to co-localize in siVAE feature embedding space.

## RESULTS - Gene hubs can be identified without explicit gene network inference

Our observation that hub genes in a community are treated differently by siVAE led us to hypothesize that we may be able to identify hub genes from a trained siVAE model without inferring a GCN. Hub genes are often identified after GCN inference because they play essential roles both in terms of the structure of the network and the genome itself, and are often targets of genetic variants associated with disease^62,63^. We reasoned that because hub genes are connected to so many other genes, siVAE is more likely to store the expression patterns of hub genes in the compressed representation (latent embedding) for use in reconstructing the rest of the gene expression patterns. We therefore hypothesized that we could identify hub genes as those genes that are well reconstructed by a trained siVAE model, because if siVAE captures variation in hub gene expression in the cell embedding space, it should also reconstruct the hub gene expression more accurately than other genes. We therefore used gene-specific reconstruction accuracy in the siVAE model as GCN -free measure of degree centrality. As a ground truth measure of gene centrality, we calculated each gene’s individual ability to predict the expression levels of every other gene in the genome (see Methods), reasoning that a ‘hub’ gene should be predictive of many other genes in the network.

**Figure 5a** compares siVAE’s estimate of gene centrality with gene centrality calculated on GCNs inferred using a number of existing GCN inference algorithms (see Methods). Overall, siVAE has the highest correlation between its predicted gene centrality and the ground truth centrality (Spearman *ρ*=0.90, P=2.2×10^-16^), significantly larger than other approaches (median Spearman *ρ*=0.36, P=9. x10^-11^). When identifying hub genes in a network, it is typical to focus on the genes with highest predicted centrality. We found that the top 20 genes with highest predicted degree centrality for siVAE has mean ground truth degree centrality of 0.092. This compares favorably to the GCN inference methods, for whom the median of mean ground truth degree centrality is 0.074 for the top 20 most central genes identified by the GCN inference methods. **Figure 5b** illustrates the cumulative ground truth degree centrality of the top predicted hubs according to each method, and siVAE consistently selects genes with the largest cumulative degree centrality of all tested methods. These results in total suggest that using siVAE, we can identify high degree centrality genes more accurately than the more classic approach of first inferring a gene co-expression network before identifying high degree centrality genes.

**Figure 5:**
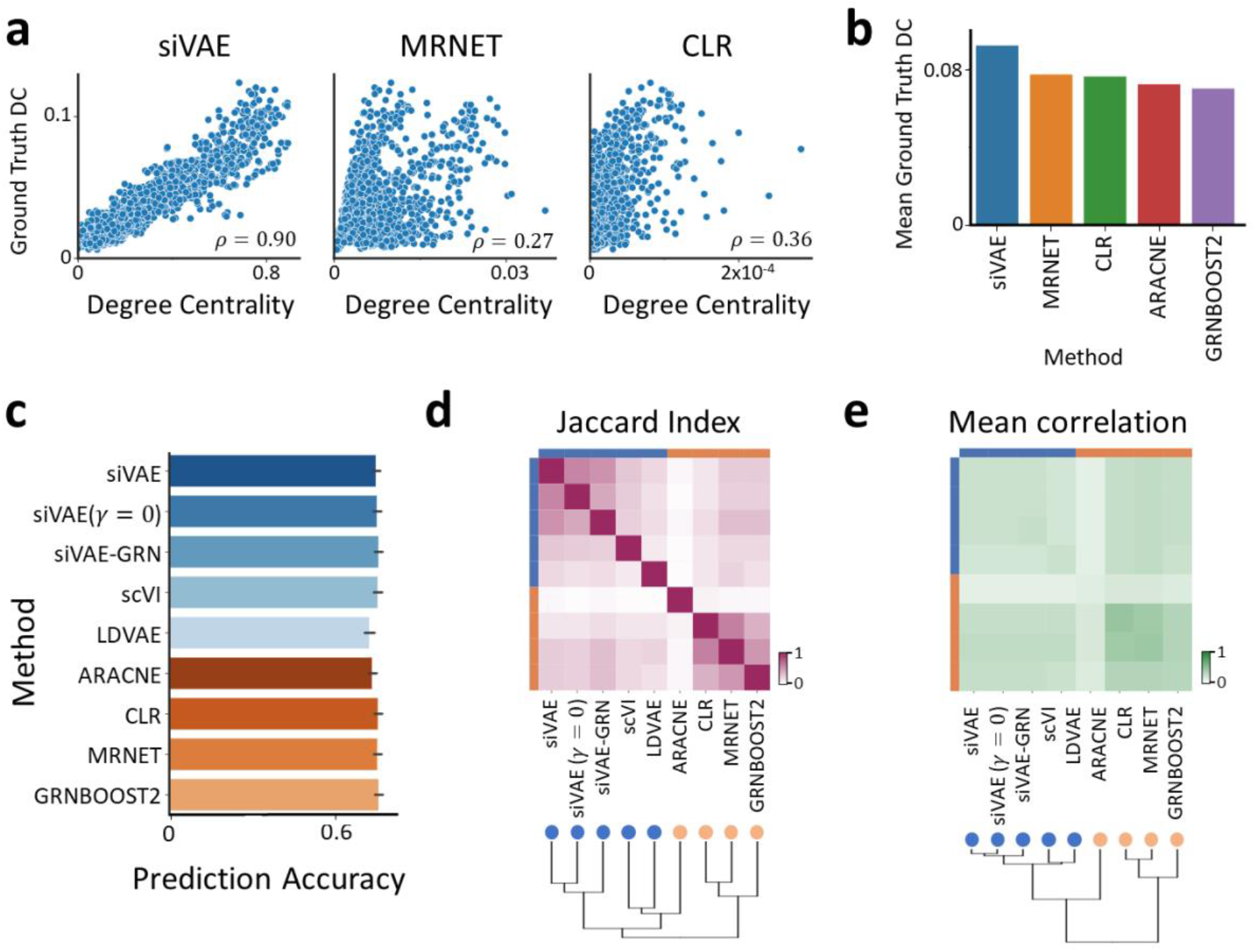
siVAE can be used to gain insight into gene co-expression network structure without explicit network inference. (**a**) Scatterplot showing the correlation between ground truth degree centrality and predicted degree centrality, based on using siVAE training performance, or by computing node degree when a network is inferred using the MRNET or CLR algorithms. (**b**) Average true degree centrality of the top 50 genes predicted to have highest degree centrality across different methods. (**c**) Bar plot indicating the prediction accuracy (% variance explained) of the neighborhood gene sets when predicting each query gene, averaged over the top 152 query genes with highest predicted degree centrality across all tested methods in the fetal liver atlas dataset. Blue bars denote methods based on dimensionality reduction, while orange bars indicate methods based on explicit gene regulatory network inference. (**d**) Heatmap indicating the pairwise Jaccard index (overlap) between neighborhood genes identified by pairs of methods. (**e)** Heatmap indicating the mean pairwise correlation in expression between pairs of methods with respect to their neighborhood genes. Each box indicates average of the pairwise correlation matrix where the columns and rows correspond to neighborhood genes identified by two methods.

## RESULTS - Systematic differences in gene neighbors identified by dimensionality reduction and network inference methods

Finally, we explored the extent to which we could identify neighboring genes that share an edge in a GCN, without having to infer GCNs explicitly. Gene neighbors tend to share similar function^64^, interact with one another^65^ and/or belong to the same gene community^66^. Identification of gene neighbors therefore aids in identifying co-functional genes in the cell.

Here we hypothesized that we could identify gene neighbors directly using a trained siVAE model, instead of having to first infer an explicit GCN. GCN inference methods typically output edge weights between pairs of nodes in the network, where larger weights correspond to a greater chance the two nodes share an edge in the underlying GCN. For siVAE, we generate edge weights in two ways: (1) siVAE-Euc, the Euclidean distance of the two genes in the feature embedding space, where smaller distances correspond to closer proximity, and (2) siVAE-GCN, where we first sample a new scRNA-seq dataset from siVAE that matches the size of the training data, then run a GCN inference method (ARACNE, MRNET, CLR, and GRNBOOST2) on the sampled scRNA-seq dataset to calculate edge weights between genes. To quantitatively evaluate the accuracy of neighbor identification using each method, we measured the percentage variance explained of a given query gene when predicted by the expression levels of the nearest 20 genes ranked by edge weight to the query gene (see Methods). In our evaluations, we only consider the 152 query genes which were predicted to have high degree centrality across all tested methods (see Methods).

Overall, most methods identified neighbors that were equally predictive of the 152 query genes’ expression levels (**Fig. 5c**). Excluding LDVAE and ARACNE, the median % variance explained for each method was 79.9% ± 0.84 s.d. **Supplementary Fig. S14** illustrates that excluding LDVAE and ARACNE, the pairwise difference in % variance explained between methods is only 0.013% on average. Notably, we observed lower % variance explained for LDVAE and ARACNE (on average, 77.2% variance explained, and 78.3% variance explained, respectively). The poorer results of LDVAE are consistent with our classification performance results above.

When considering the overlap in neighbors selected by different methods, it is striking how the dimensionality reduction methods cluster strongly (scVI, siVAE, LDVAE) and the GCN inference-based methods cluster strongly as well, with markedly less overlap between these two groups (**Fig. 5d)**. This is surprising in part because the neighborhood sets are all approximately of the same predictive performance (**Fig. 5c**), suggesting the DR methods are systematically identifying different neighbors that are as equally co-expressed as the neighbor set identified by the GCN methods. In particular, consider that siVAE-GCN involves identifying gene neighbors using the GCN inference methodology, but just applied to a siVAE-generated dataset (instead of the original training dataset). **Figure 5c** illustrates that under the siVAE-GCN neighborhood identification framework, the neighborhood genes are still much more similar to siVAE than to the GCN inference methods, suggesting the unique neighborhood identified by the DR methods is a property of the co-expression patterns that DR methods learn, and not due to the way in which neighborhood genes are identified. The poor overlap between the DR and GCN methods also holds true if we consider the average pairwise correlation in expression between neighbor sets, instead of considering overlap of genes (**Fig. 5e**). More specifically, the GCN-defined neighbor sets had higher average Pearson correlation amongst themselves (average Pearson correlation = 0.67, excluding ARACNE) compared to the average Pearson correlation coefficient among the neural net-based neighbor sets (average Pearson correlation = 0.46). There was also low average correlation between DR and GCN neighbor sets (average Pearson correlation = 0.39). Our results therefore suggest that since GCN- and dimensionality reduction-identified neighbor sets are systematically different but approximately equally predictive of neighboring genes, then both approaches should be used to find co-expressed genes in a network.

## RESULTS - Co-regulation of mitochondrial genes in iPSCs linked to neuron differentiation efficiency

We next wondered whether we could leverage siVAE to compare GCN structure across multiple cell populations and associate changes in network structure with cell population phenotypes. Because robust single-GCN inference is already challenging, there has not been extensive work into approaches to comparing multiple GCNs^67–69^. As introduced earlier, the NeurDiff dataset includes scRNA-seq data collected across 215 iPSC cell lines profiled before differentiation (at 11 days), as well as after initiation of differentiation into neurons (day 30 and 52). By computing the fraction of sequenced cells at day 52 that were neuronal, each cell line has an estimate as to how efficiently they could be differentiated into neurons. Efficiencies were found to be highly reproducible and potentially associated with expression profile of the pluripotent cells at Day 11^37^. Thus, we focused on the identification of GCN structure in iPSCs (at day 11) that were associated with neuronal differentiation efficiency (measured at day 52) (**Fig. 6a**). The pluripotent cells sequenced at day 11 were further divided into two cell types based on whether they were proliferating (P_FPP) or non-proliferating (FPP) mid brain floor plate progenitors, and we performed GCN analysis on P_FPP and FPP cells separately.

**Figure 6:**
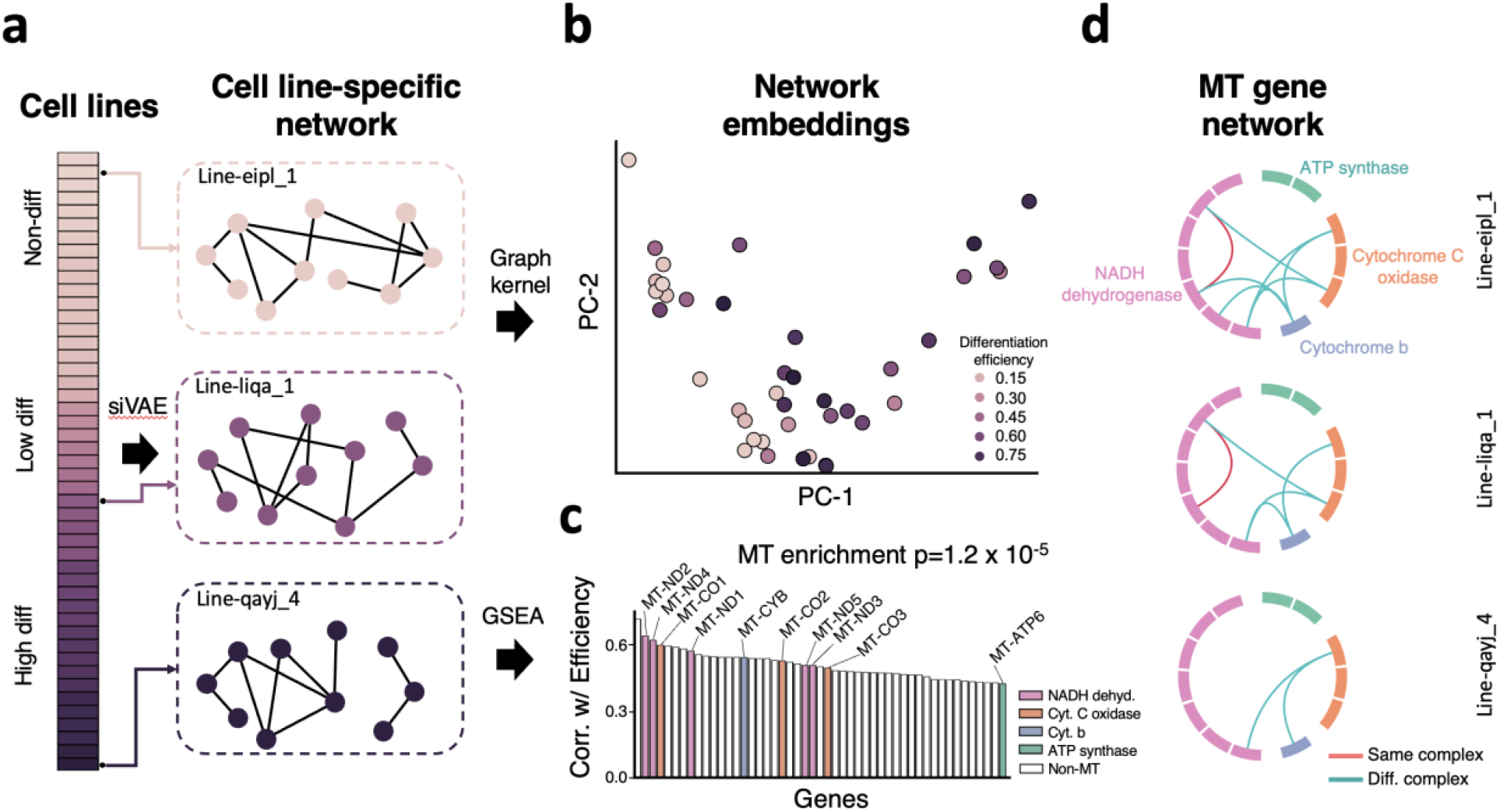
Connectivity between mitochondrial genes decrease for iPSC cell lines with higher differentiation efficiency. (a) Schematic for generating cell line network embeddings. Each of the stacked bar indicates an iPSC cell line whose gene network is inferred with siVAE. (b) Scatterplot of cell line embeddings. Each dot indicates cell line in the cell line-to-cell line matrix generated from graph kernel trained on siVAE inferred gene network. (c) Bar plot showing Spearman correlation between degree centrality and differentiation efficiency. Each bar indicates a gene, and the colored bars belong to mitochondrial genes. Each color represents mitochondrial complex the gene is coding for. P-value (d) Visualization of mitochondrial genes in siVAE inferred gene network for three selected cell lines. Only 7 connections that are significantly less likely to be present in cell lines with higher efficiency are being displayed.

We first trained siVAE on each of the 41 iPSC lines to yield 41 gene embedding spaces. We then calculated a pairwise distance between every pair of iPSC lines using a graph kernel on the gene embedding spaces (see Methods). **Figure 6b** visualizes these distances between iPSC lines using PCA. Surprisingly, we observed separation of cell lines according to their neuronal differentiation efficiency along PC-1, when visualizing all lines together (Fig. 6b), as well as when visualizing only iPSC lines that showed any differentiation (**Supplementary Fig. 12**). This result was unexpected because the graph kernel-based visualization (**Fig. 6b**) does not use any information about neuronal differentiation efficiency to guide visualization. The fact that the iPSC lines still separate by efficiency suggests that the major differences in network structure between iPSC lines is strongly associated with differentiation efficiency. We confirmed on both the two progenitor cell types (P_FPP and FPP) that differentiation efficiency explains separation in the cell line embeddings (**Supplementary Fig. 15**), but we focused on FPP for downstream analysis because of larger dataset size.

To determine the genes whose varying structure is most responsible for explaining variation in **Figure 6b**, we computed siVAE degree centrality for each gene and for each iPSC line, and used gene set enrichment analysis (GSEA) to identify which gene sets and pathways whose degree centrality correlated with differentiation efficiency. Overall, we found 125 genes’ degree centrality was significantly correlated with differentiation efficiency (**Supplementary Table 12**). Of these 125 genes, 10 of them are from the mitochondrial (MT) genome. We found that the degree centrality of all 10 MT genes were significantly correlated with differentiation efficiency (GSEA, adjusted P=1.2×10^-5^) (**Fig. 6c**). Individually, mitochondrial genes had high correlation between their degree centrality and efficiency, with median Spearman correlation of 0.53 and P-value of 2.3×10^-5^.

A common explanation for change in connectivity for a single gene in a GCN is a change in overall expression; genes that are turned off cannot covary with other genes, for example. To rule out this trivial explanation and focus on genes whose degree centrality is correlated with differentiation efficiency independent of changes in mean expression, we identified genes whose mean expression is correlated with differentiation efficiency. None of the individual mitochondrial genes’ mean expression levels were significantly correlated with differentiation efficiency (median Spearman *ρ*=0.016, median adjusted P=0.75, **Supplementary Fig. 16**). In contrast, 88 of the 115 non-MT genes whose degree centrality correlated with efficiency also demonstrated high correlation between efficiency and mean gene expression (median Spearman *ρ*=0.21, median adjusted P=6.0×10^-6^). The correlation between MT genes’ degree centrality and differentiation efficiency is therefore not explained by changes in mean expression of MT genes.

Next, we examined the specific changes in MT genes’ network connectivity that were driving their correlation between degree centrality and efficiency. Based on the GCN inferred from siVAE gene embeddings, the number of edges between MT genes and non-MT genes were consistently low across all cell lines with an average of 0.65 edges per cell line and showed insignificant correlation with efficiency (Spearman *ρ*=0.20, P=0.14) (**Supplementary Fig. S17)**. Connectivity between MT and non-MT genes therefore does not explain the variation in MT gene connectivity across lines. Instead, the correlation between MT gene centrality and differentiation efficiency is driven by changes in connectivity within the MT genes with an average of 73 edges per cell line and high correlation with efficiency (Spearman *ρ*=-0.59, P=1.5 x10^-6^).

The mitochondrial genes encode for subunits of mitochondrial complexes. Given our results above suggest that connectivity between MT genes is correlated with differentiation efficiency, we sought to distinguish if it were connections between MT genes in the same or different complexes that are correlated with differentiation efficiency. We performed a Wilcox rank sum test on every pair of MT genes to test whether edges between or within MT complexes are correlated with differentiation. We identified seven MT edges correlated with differentiation efficiency in total and of the seven, six of them connect pairs of MT genes from different complexes. Of these six edges, three connect NADH dehydrogenase to cytochrome C oxidase, two edges connect NADH dehydrogenase to cytochrome b, and one edge connects cytochrome C oxidase to cytochrome b. This suggests that coregulation of distinct MT complexes is an important indicator of differentiation efficiency of these iPSC. Given the overall importance of MT genes and the potential role they play in differentiation efficiency, we then looked for genetic variants in MT genes that are associated with differentiation efficiency. Unfortunately, we were unable to identify genetic variants in the MT genome that were significantly correlated with differentiation efficiency (**Supplementary Note 2**).

## Discussion

Through the development of siVAE, we have mitigated one of the primary limitations of the interpretation of VAEs: the slow execution time of neural network feature attribution methods when the number of input features and embedding dimensions of the cell embedding space are both large. Single cell atlases are ever-increasing in size due to the dropping cost of single cell sequencing^70^. Also, there is rapidly increasing interest and development of multi-modal single cell assays such as SNARE-Seq^71^, ECCITE-Seq^72^, and SHARE-Seq^73^ that measure multiple data modalities (RNA, ATAC) simultaneously and are yielding single cell measurements with up to hundreds of thousands of input features, which will then demand large cell embedding spaces to accurately capture covariation in input features. As such, we expect the importance of scalable, interpretable VAEs will continue to grow.

Our analysis has also demonstrated how interpretation of cell embedding spaces can lead to insight into the gene regulatory networks underlying the cell population siVAE is trained on. In addition to showing how co-expressed groups of genes can be identified, we also showed how we can identify hub genes, without inferring an explicit GCN. This is useful because GCN inference continues to be a highly challenging task, even in the era of large numbers of cells sequenced from single cell assays^74^. Furthermore, siVAE can also find groups of co-regulated genes that are not readily identified by GCN methods, suggesting the two approaches to identifying co-regulated gene sets are complementary in their findings. By comparing VAEs trained on different cell populations, it is likely possible to identify differential co-expression patterns between cell populations, also without explicit GCN inference. Finally, we demonstrate siVAE identifies genes with high degree centrality more accurately than ranking genes by explicit node degree in a gene network, suggesting siVAE can be used to find central genes in the genome.

A surprising observation we made was that the set of neighbors of a given gene with respect to the underlying GCN is systematically different between explicit GCN inference methods and the dimensionality reduction methods. This even holds true when a trained siVAE model was used to sample expression data that was then sent as input into a classic GCN inference method; in this scenario, the resulting GCN yielded neighbors that were similar to those identified directly from the DR methods. Our experiments further showed that both neighborhood sets are equally coexpressed with the query gene, suggesting at the least that accurate GCN construction should leverage both of these types of approaches to identifying gene neighbors. One possible explanation is that DR methods can learn to combine many genes into a single embedding dimension, whereas explicit GCN inference methods ultimately represent co-expression patterns as individual edges between only pairs of genes, and therefore are more limited in their capacity to represent higher order co-expression patterns.

Previous studies have established the importance of mitochondria in reprogramming, maintenance of pluripotency, and differentiation through their functional role in energy production ^75–77^. With respect to gene regulation, key mitochondrial transcription factors influence iPSC’s ability for differentiation^78–82^. The mean expression levels of established pluripotency markers such as SOX2, Oct4, Nanog, Klf4, and c-MYC^83^ are correlated with differentiation efficiency^37^. Also, transcription factors associated with mitochondrial biogenesis (TFAM, POLG1, and POLG2)^77,78^ are necessary for successful differentiation. Our results showing coregulation of MT complexes as an indicator of differentiation efficiency are complementary in that there are few studies identified additional downstream genes^84,85^ associated with differentiation efficiency; prior work focused on identifying genes whose mean expression was correlated with differentiation efficiency, and did not identify MT genes^37^.

There has also been recent work studying the impact of mitochondrial heteroplasmy on iPSC differentiation potential. Several studies now suggest mtDNA integrity as mandatory iPSC selection criteria^86–88^. Heteroplasmy of several mutations have been linked to iPSC’s ability to differentiate^87–90^. Manipulating MT heteroplasmy through insertion of wild type mtDNA has been shown to revert diseased iPSC state and improve pluripotency^91^. Correlation of heteroplasmy with co-expression of MT complexes is an interesting avenue to pursue to determine whether heteroplasmy may be a cause of de-correlation of MT complexes.

While we have chosen the classic VAE framework upon which to build siVAE, our approach to introducing a feature-wise decoder and interpretability term is generalizable and can be applied to other general extensions of VAE, such as VAE-GANs, β-VAE among others ^92,93^. With respect to genomic data modalities such as epigenomics, miRNA, and scRNA-seq, methods such as SCALE^94^, RE-VAE^24^, methCancer-gen^25^, VAEMDA^27^, scMVAE^26^, scVI^20^, Dr.VAE^32^, scGen^33^, and Dhaka^29^ could also benefit from similar interpretability terms such as that used for siVAE. Many of these methods specifically focus on analysis beyond visualization^20^,^28^ such as trajectory inference^29^,^95^, data imputation^30^, and perturbation response prediction^31–33^. An additional interpretability term could enable identification of key input features in each task (e.g. which set of regulatory genes are tied to differentiation progression, which input genes are used to impute missing genes, and which sets of genes are affected by drug perturbation), which is a crucial step for validation and downstream application of these methods. Recently, there has been great interest in integrative analysis of multi-modal data resulting from sequencing technologies that measures two modalities such as SNARE-Seq^71^ measuring gene expression and chromatin accessibility as well as CITE-Seq^96^ measuring gene expression and protein expression. Even for methods that modifies VAE to jointly model two modalities (totalVI^97^, multiVI^98^, scMVP^99^, BABEL^100^, and Cobolt^101^), an interpretability term could be applied to understand how individual features of each modality relate to features of another modality such as linking enhancers based on chromatin accessibility to its target genes.

A related set of approaches to increasing interpretability of generative models focuses on disentanglement learning. In particular, methods such as InfoGAN^102^, FactorVAE^48^, DirVAE^103^ and others^93,104,105^ modify generative models such as the VAE to achieve disentangled representation by encouraging the individual cell dimensions to be statistically independent. They show that independence between cell dimensions oftentimes leads to more correspondence between individual cell dimensions and tangible factors such as width and rotation of digits for MNIST. However, we do not consider these model variants here because they do not provide contributions of individual features to cell dimensions. These approaches still require users to manually draw samples of points from the cell embedding space, reconstruct the input features from the cell dimensions, then use human intervention to manually inspect how variation across specific dimensions might correspond to human-interpretable factors of variation. However, the regularization terms that encourages disentanglement between the cell dimensions may be applied to siVAE. This would help remove the entanglement between cell dimensions such as the overlapping outlines of digits in siVAE loadings for the MNIST dataset.

## ACKNOWLEDGEMENTS

This publication was made possible by an NIGMS funded predoctoral fellowship to YC (T32 GM007377). GQ was supported by NSF CAREER award 1846559. This project has been made possible in part by grant number 2019-002429 from the Chan Zuckerberg Foundation. This work was supported by the National Institute of Child Health and Human Development P50 HD103526.

## METHODS

### Model notation

We denote vectors as lower case, bold letters (e.g. z). Matrices are upper case letters with two subscript indices (e.g. *X_c,f_*). Constants are upper case letters with no subscripts (e.g. *L*).

### Generative process of VAEs

siVAE is an extension of a classic variational autoencoder. Here we briefly review the generative process assumed by a standard VAE with *L* hidden layers in the decoder, and in which the hidden units of the last layer of the decoder are linearly transformed into the predicted mean of the Gaussian distribution over the observed data:

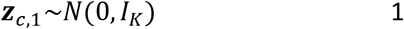

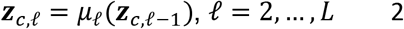

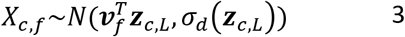

*X_c,f_* is the input observed value for feature (e.g. gene) *f* and cell *c* (centered and scaled across all cells), where we assume there are *F* features and *C* cells in the training data. *z*_*c*,1_ is the embedding of cell *c* in the (latent) cell embedding space of the VAE, while *z_c,ℓ_* for *ℓ* > 1 represent the activations of the hidden layer *ℓ* of the decoder for cell *c. **v**_f_* is the vector of incoming weights to the predicted mean of the output node *f* of the VAE, while *σ_d_*(·) is a one-layer function that predicts a non-negative scalar value representing variance. *I_K_* is the identity matrix of rank *K*. *μ*_1_(·), …,*μ_L_*(·) represent the parameterized activation functions of hidden layers 1, …,*L* of the cellwise decoder, respectively.

### Generative process of siVAE

The key idea behind siVAE is that we jointly infer cell-wise and feature-wise state spaces, and through regularization, loosely enforce correspondence between the cell and feature dimensions. Here, correspondence means variation in dimension *k* in the cell embedding space corresponds to observed variation in each feature *f* that is proportional to feature *f*’s embedding coordinate in dimension *k*. Through correspondence, the feature embedding coordinates (‘siVAE loadings’) become analogous to factor loadings, and the cell embedding coordinates (‘siVAE scores’) become analogous to the factor scores of PCA. In siVAE, the feature and cell embeddings are sampled from different latent spaces and projected to higher dimensions through separate decoders, before combining to produce the means of the Gaussians (**Figure 1a**). The generative process assumed by siVAE is shown below:

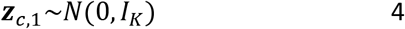

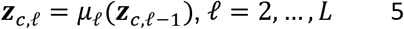

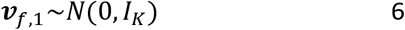

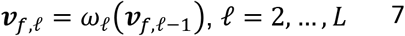

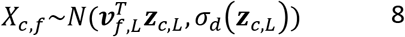

Here ***z**_c,L_*, *μ_ℓ_*(·), *I_K_* and *σ*(·) are defined as above for VAEs. ***ν***_*f*,1_ is the latent embedding of feature *f* in the feature embedding space of siVAE, while ***ν**_g,ℓ_* for *ℓ* > 1 represent the activations of hidden layer *ℓ* of the feature-wise decoder for feature *f*. *ω*_1_(·), …,*ω_L_*(·) represent the activation functions of hidden layers 1, …,*L* of the feature-wise decoder, respectively.

Comparing Equations 3 to Equations 6–8 illustrate that siVAE turns the last layer of weights leading to the Gaussian mean of the VAE into a non-linear transformation of the latent variables ***ν***_*f*,1_. siVAE can therefore be viewed as putting a prior over a single (last) layer of weights in the VAE. The matrix *V* = [*ν*_1,1_,…, *ν*_*F*,1_]^T^ encodes the siVAE loadings, while the matrix *Z* = [***z***_1,1_, … ,*z*_*C*,1_] encodes the siVAE scores. Note that we can compute siVAE loadings and scores of other hidden layers *ℓ* as well, but in this paper, we focus on the latent space (*ℓ* = 1).

### Inference and training

We employ variational inference via a pair of encoder networks, *ψ*(*X_;,f_*) for features and *ϕ*(*X*_*c*,:_) for cells, in a manner analogous to variational inference applied to VAEs. Note the input for the two encoders is different: *X_:,f_* is a vector of observations for a single feature *f* across all training cells, whereas *X*_*c*,:_ is a vector of observations for a single cell *c* across all features. Our approximate posterior 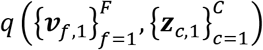 factors as follows:

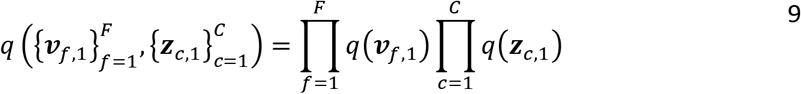

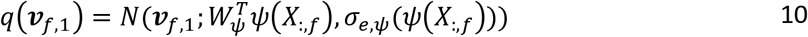

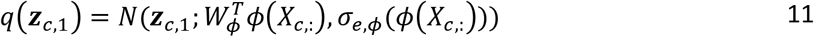

We perform variational inference and learning by maximizing the expected lower bound function *ℓ*_SIVAE_, where 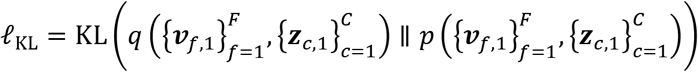.

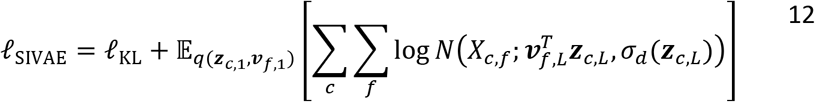

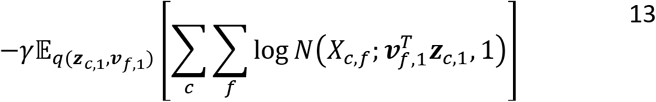

### Interpretability term

The right-hand side of Equation 12 is analogous to the KL divergence and reconstruction loss terms of the original VAE lower bound function. The term in Equation 13, which we call the interpretability term, encourages the individual embedding dimensions of ***ν***_*f*,1_ and ***z***_*c*,1_ to correspond to each other, by encouraging the linear products between ***ν***_*f*,1_ and ***z***_*c*,1_ to approximate *X_c,f_*. In our experiments, we set the penalty term *γ* = 0.05 to make the effect of the interpretability term small on the overall loss function.

### Reducing dimensionality of input for feature-wise encoder-decoder

The size of input *X_:,f_* for feature-wise encoder-decoder increases with *C*, number of cells. To avoid the computational expenses of models with potentially millions of input nodes, we reduce the dimensionality of the input from *C* to *C_red_* through either downsampling or PCA. For downsampled input, we randomly sample *C_red_* cells while maintaining the ratio between the cell types which accounts for redundancy of information between cell types with similar gene expression patterns. For PCA input, we performed PCA without whitening on *X^T^*, *G x C* matrix and retained first *C_red_* principal components resulting in *X’^T^*, *G x C_red_* score matrix. The preservation of linear covariation with PCA is analogous to common usage of PCA before t-SNE or UMAP. In **Supplementary Fig. S16**, we show training the feature-wise encoder-decoder with downsampled and PCA inputs both results in loss and clustering accuracy score comparable to that of model trained with the full data.

### Training procedure for siVAE

We use a three-step training procedure to improve inference and learning, described in more detail in the Supplementary Materials:

- **Pre-train cell-wise encoder and decoder**. We first train the cell-wise encoder and decoder, similar to how a classic VAE is trained, by optimizing the Equation 12 component of *ℓ*_SIVAE_ with respect to {*μ_ℓ_, σ_d_, ϕ, σ_e,ϕ_, W_ϕ_*}, and by treating the variables *ν_f,L_* as parameters to optimize to estimate 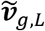. The input to the cell-wise decoder are the cell-wise data points *X*_*c*,:_, and the output are the same data points *X*_*c*,:_.
- **Pre-train feature-wise encoder and decoder**. We next train the parameters associated with the feature-wise encoder and decoder, namely {*ω_ℓ_,ψ,σ_e,ψ_ψ,W_ψ_*}, by training a VAE whose inputs are the data features *X*_:,*f*_, outputs are 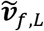 learned from the previous step, and whose encoder is defined by {*ψ, σ_e,ψ_, W_ψ_*}, and decoder parameterized by *ω_ℓ_*, for *ℓ* = 1,…,*L*–1.
- **Train siVAE**. We finally train all model parameters {*μ_ℓ_,σ_d_,ϕ,σ_eϕ_,W_ϕ_,ω_ℓ_,ψ,σ_eψ_,W_ψ_*} jointly by optimizing the full *ℓ*_SIVAE_ from Equation 12,13.

### siVAE and VAE network design

For our experiments, identical neural net designs were used across the feature-wise and cell-wise encoders and decoders in siVAE. The architecture of the VAEs we compared against were matched to the architecture of the cell-wise encoder/decoders of siVAE. For MNIST and Fashion-MNIST, we set the architecture of the encoder to two hidden layers of sizes 512 and 128, and the decoder to two hidden layers of sizes 128 and 512. For all other datasets except the LargeBrainAtlas dataset, we set the architecture of the encoder to three hidden layers of sizes 1024, 512 and 128, and the decoder to three hidden layers of sizes 128, 512 and 1024. LargeBrainDataset, we trained an encoder with three hidden layers of sizes 2048, 1024, and 512, and the decoder with three hidden layers of sizes 512, 1024, and 2048. We used a latent embedding layer with size varying between 2, 5, 10 and 20 nodes for all imaging datasets. For the fetal liver atlas, we set the latent embedding layer size to be 2 for visualization and 64 for all other cases. In the timing experiment, we varied the latent embedding layer size between 20, 128 and 512 for the LargeBrainAtlas dataset, while setting the latent embedding layer size at 2 for the scATAC-Seq dataset. For the NeurDiff dataset, we set the size of the latent embedding layers to be 32. Additional implementation details as well as a table containing the above information on network design can be found in the Supplementary Materials (**Supplementary Table 1**).

### siVAE and VAE model selection

We set model hyperparameters and optimization parameters by performing a hyperparameter search for the model with lowest total loss on the held-out data. For each model, we used the Adam optimizer for training, with a learning rate of either 0.0001, 0.001, or 0.01. We considered L2 regularization with scale factor *λ* of either 0.001 or 0.01. For imaging datasets, we set the number of embedding dimensions to 20. For genomic datasets, we used two embedding dimensions for models that were used for visualization, and otherwise considered sizes of 16, 32 and 64 for all other analyses.

### siVAE model variants

To explore the role of different design choices of siVAE, we created several variants of the siVAE model described above. siVAE (*γ* = 0) removes the interpretability term in Equation 13 (by default, *γ* = 0.05). For comparison against LDVAE, whose decoder network ultimately predicts the parameters for negative binomial distributions instead of the parameters of a Gaussian distribution as implemented in siVAE and VAE, we also implemented both siVAE (NB) that predicts the parameters of a negative binomial distribution and VAE (linear) that is an identical implementation of LDVAE. siVAE (NB) is formulated as follows, where *l_μ_, l_σ_* parametrize the prior for scaling factor and are set to empirical mean and variance of the observed data:

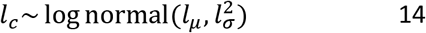

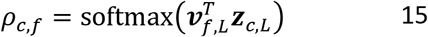

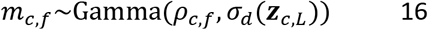

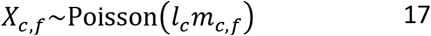

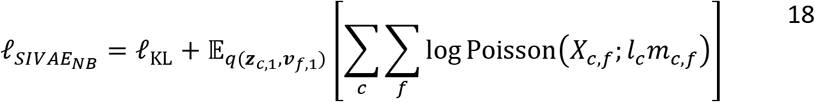

VAE (linear) is identical to siVAE (NB) except 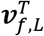 is replaced by *ϕ_f_*, an estimated parameter that matches the length of ***z**_c,L_*, thereby removing thefeature-wise encoder-decoder from the model. Finally, we implemented siVAE (linear), where the mean of the distribution over *X_c,f_* is directly predicted from linear multiplication of cell and feature embeddings. The reconstruction loss term corresponds to interpretability term, eliminating need for the latter.

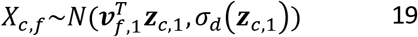

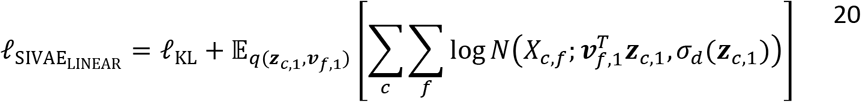

Batch correction is natively implemented in siVAE with a similar approach used in scVI^20^. *s_c_* is a vector of length *b* whose individual element is either a continuous feature or one-hot encoding of categorical feature. The batch vector is concatenated to the input of the cell-wise enc-dec as well as the cell embedding to minimize the amount of batch effect captured in the cell embedding.

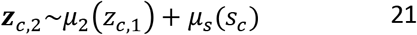

Additionally in the interpretability regularization term, we add weight *j_f_*, a vector of length *b*, that accounts for batch effect absent in the linear reconstruction in cell embedding.

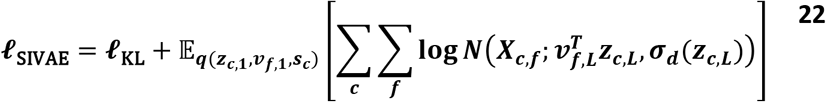

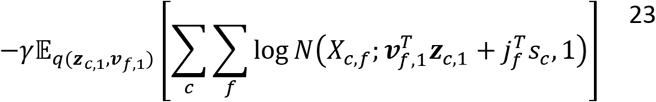

### siVAE model availability

siVAE is implemented as a Python package and is available from PyPi (https://pypi.org/project/siVAE/).

### LDVAE and scVI

We used LDVAE^15^ and scVI^20^ implemented in SCANPY^106^ package available from PyPi. The architecture of the model was set to match that of the cell-wise encoder-decoder of siVAE, including the number of dimensions of the cell embedding space and the number of hidden layers, as well as the number of hidden nodes. Model optimization was performed by varying learning rate between 1e-2, 1e-3, and 1e-4 while the rest of the parameters were set to default.

### Feature attribution methods

Two separate Python packages were used to compute neural network feature attributions in our experiments. We used the DeepExplain Python package that implemented all feature attribution methods (Saliency Maps, Grad*Int, DeepLIFT, IntGrad, Shapley Value) included in our experiments in reverse-mode^107^. We used the tensorflow-forward- ad Python package for computing Saliency Maps and Grad*Int in forward-mode^108^. In both cases, the package applies feature attribution between the target nodes and input nodes. For application of feature attributions on the decoder, the target nodes and input nodes were set to be the nodes of the output layer and latent embedding layer, respectively, for the cell-wise decoder. For application of feature attributions on the encoder, the target nodes and input nodes were set to be the nodes of the latent embedding layer and input layer, respectively, for the cell-wise encoder. By default, the DeepExplain package summarizes the attribution across all target nodes, so binary masks corresponding to a single target node were used per target node. Similarly, the tensorflowforward-ad package summarizes attribution across all input nodes, so binary masks corresponding to a single input node were used per input node. Integrated Gradients and DeepLIFT require an additional parameter of input baseline, which represents a default null value that input values can be referenced against. We set this value to 0 equaling the mean value of gene expression after preprocessing.

For Gene Relevance, we used the published R package^48^. The method required the latent embeddings learned from siVAE as well as the raw count data corresponding to the embeddings. We also varied the number of neighborhoods (10, 100, 1000, and default).

### Feature embeddings for feature attribution methods and Gene Relevance

All feature attribution methods tested here can output feature importance scores ***s**_f,c_* that represents a vector of contributions of feature *f* to all cell dimension *d* for cell *c*. The Gene Relevance method^48^ outputs partial derivatives in the same format. In contrast, siVAE loadings ***ν***_*f*,1_ represents a vector of contributions of feature *f* to all cell dimensions, summarized over all cells. To compare feature attribution methods to siVAE, we therefore need a procedure for converting the per-cell attributions ***s**_f,c_* into a set of overall feature attributions ***u**_f_* for each feature f with respect to all cell dimensions and that summarize across all cells, analogous to siVAE’s loadings ***v***_*f*,1_. To do so, we first construct a matrix *S_d,f,c_*, containing all feature attributions for cell dimension *d*, cell *c* and feature *f*. For each cell dimension *d*, we apply PCA to the 2D matrix *S*_*d*,:,:_. to extract the first principal component’s loadings ***u***_:,*d*_, a vector of length *F* that contains the contribution of each input feature *f* to cell dimension *d*. Repeating this process for each cell dimensions then concatenate the resulting vector results in matrix ***U**_f,d_*, whose rows ***U***_*f*,:_ are analogous to siVAE’s *ν*_*f*,1_. Finally, we calculated Spearman correlation with two-sided test between the feature embeddings inferred through different approaches per dimension and reported the median values.

### Datasets

A table summarizing the following datasets can be found in **Supplementary Table 2.**

### Fetal liver atlas dataset processing

We obtained the fetal liver atlas^34^ from ArrayExpress with accession code E-MTAB-7407 (https://www.ebi.ac.uk/arrayexpress/experiments/E-MTAB-7407/) on 2020/06/10, in processed form. We normalized the count matrix to TP10K, then performed feature selection by retaining the top 2,000 highly variable genes, yielding 177,376 cells and 2,000 genes. We then downsampled the number of cells to 100,000 cells, while preserving the proportion of cells from each cell type. Genes were individually centered and scaled to unit variance. For visualization of the feature embeddings for the liver fetal atlas in **Figure 3b**, we obtained marker genes for four cell types (hepatocytes, Kupffer cells, NK/NKT cells, and MHC II positive B cells) that were available in the MsigDB database^109^ (downloaded from http://www.gsea-msigdb.org/gsea/msigdb/collections.jsp#C8 on 2020/02/08). To account for the multiple subtype labels in the fetal liver dataset matching to a single cell type in marker gene set, we allowed many-to-one mapping by grouping multiple cell type labels in the dataset that corresponded to one of the cell types in marker gene database. The exact groupings are shown in **Supplementary Table 3.** We only visualized the cells with known marker genes, and genes that belonged to more than one marker gene set (shared across cell types as a marker) were discarded.

### MNIST and Fashion-MNIST dataset processing

We obtained both datasets from the TensorFlow datasets web page on 2020/02/20. Images were flattened and centered and scaled to unit variance per feature across all images before input into the models.

### CIFAR-10 dataset processing

We obtained CIFAR-10 from the TensorFlow datasets web page on 2020/02/20. We then subsampled the image classes to only the airplane and ship classes because other image classes require convolutional layers to achieve good classification performance, but here our goal was to benchmark VAEs. Images were flattened and centered and scaled to unit variance per feature across all images before input into the models. Color channels were concatenated and flattened.

### LargeBrainAtlas dataset processing

We obtained the 1.3 Million Brain Cells dataset referred to as “LargeBrainAtlas” from the 10x Genomics website (https://support.10xgenomics.com/single-cell-gene-expression/software/pipelines/latest/advanced/h5_matrices) on 2020/04/28. We normalized the count matrix to TP10K, then retained all genes. After, genes were individually centered and scaled to unit variance.

### BrainCortex dataset processing

We obtained the BrainCortex dataset (GSE126074) from (https://www.ncbi.nlm.nih.gov/geo/query/acc.cgi?acc=GSE126074) on 2020/12/01. We performed quality control based on TSS enrichment and nucleosome signal which filtered the dataset down to 244,544 features.

### NeurDiff dataset processing

We obtained iPSC neuronal differentiation dataset referred to as “NeurDiff’ dataset from https://zenodo.org/record/4333872 accessed on 2021/12/10. We only used the gene expression matrix for day 11 (D11) with pre-normalized expression. We divided the dataset according by cell types (FPP and P_FPP) then filtered out the cell lines that contained less than 1000 cells of that cell type. The downstream preprocessing and experiments were performed per cell type. Per cell type dataset, batch correction was performed per cell line to regress out pool_id as well as cell cycle score using regress_out() function from scanpy^106^. Next, we performed feature selection by taking the union of the top 2,000 genes with highest variance in each cell line as well as the top 2,000 genes with highest variance across all cell lines. Afterwards, genes were individually centered and scaled to unit variance. The final dataset consisted of 41 cells lines (109,483 cells) with 3,362 genes for P_FPP and 27 cell liness (85,961 cells) and 3308 genes for FPP (Supplementary Table 2).

### Generation of simulated scRNA-seq datasets from a gene network

To explore the organization of genes in siVAE feature embedding space, we simulated scRNA-seq data where the correlations between genes is consistent with a specified gene co-expression network. We designed a gene co-expression network that consisted of five communities of 50 genes each, as well as an additional set of 50 isolated genes that are independently varying. Each community included a single hub gene that was connected to the other 49 genes in the community, in a hub-and-spoke model. No other genes in the community were connected to any other gene. All edge weights representing pairwise correlations between genes in the same community were set to 0.6. The adjacency matrix capturing the co-expression patterns between the 300 genes were converted to covariance matrix via the qpgraph R package^110^, using the function qpG2Sigma with parameters rho=0.6. Afterwards, we used the resulting covariance matrix as input to a multivariate Gaussian distribution and sampled 5,000 cells for training with siVAE.

### Cell type classification

The five-fold nested cross validation experiments reported in **Figure 1c** compares the performance of siVAE, VAE, and LDVAE on the fetal liver atlas dataset when matching their cell-wise encoder and decoder network designs. The number of embedding dimensions was fixed to be 2. After training using the training fold, the encoders were used to compute embeddings for the training and test datasets. We then used a k-NN *(k =* 80) classifier to predict labels of test cells based on the embeddings of the training and testing datasets. Similar five-fold nested cross validation experiments were performed on the imaging datasets (MNIST, Fashion MNIST, CIFAR-10). However, we allowed the model to individually select the number of embedding dimensions *K* from the set {2,5,10,20} using the training fold. In addition, the number of clusters, *k,* was set to 15 as imaging datasets have far fewer classes than the fetal liver atlas dataset.

### Execution time comparison experiments

We performed a series of experiments to compare siVAE training execution time against the combined execution time of VAE training and executing feature attribution methods. For Saliency Maps and Grad * Int, both forward and reverse modes were used. The majority of the feature attribution methods rely on taking the gradient of a single output nodes with respect to all input nodes using automatic differentiation in reverse-mode. For models with high number of output nodes, the operation becomes computationally infeasible. Using automatic differentiation in forward-mode allows gradient calculation of all output nodes with respect to a single input node but faces the same computational issue for models with high number of input nodes.

For the first experiment, we benchmarked using the 10x Genomics dataset. In the case of the feature attribution execution times, we extrapolated the execution time on the LargeBrainAtlas dataset from the execution time on 100,000 cells, due to time constraints and the fact that runtime of these methods should scale linearly with the number of cells. In contrast, execution times of siVAE are those measured on the full dataset. For the second experiment, we tested the effect of varying the either the number of embedding dimensions or the number of features on the execution time. As the execution time for two feature attribution methods (Integrated Gradients and Shapley Values) exceeded a realistic run time of 100 days, only the faster three methods (Saliency Maps, Grad * Int, and DeepLIFT) were used for the second experiment. For the 10x Genomics dataset, the number of embedding dimensions were set to 20, 100 and 500. Similar to the first experiment, siVAE was run on the entire set of 1.3 million cells, and the VAE+feature attribution approaches were run on 100,000 cells and then linearly interpolated to the full dataset size. For the scATAC-seq execution times, we varied the number of features by selecting the top *n* highly variable genomic regions, where *n* was set to either 28k, 120k or 240k. We used a single NVIDIA GeForce GTX1080 Ti GPU, Intel Core i5-6600K CPU, and 32 GB RAM for all experiments.

### Estimating gene centrality using siVAE

We reasoned that the expression patterns of genes with high degree centrality would be most likely to be retained by siVAE during dimensionality reduction, because those genes could be used to reconstruct the expression patterns of the many other genes connected to them. If so, then the hub genes would also be likely to be the genes whose expression patterns are reconstructed with the lowest error. We therefore define gene centrality for siVAE as the negative reconstruction error of siVAE on each individual gene during training.

### Estimating gene centrality using GCN inference methods

The GCN inference methods tested here all output pairwise weights between genes, where larger weights indicated higher confidence in a pairwise edge in the underlying GCN. We therefore measured each query gene’s degree centrality for GCN inference methods by averaging the weights between the query gene and every other gene in the network.

### Estimating the ground truth gene centrality

To compute the accuracy of siVAE-based gene centrality and GCN-based gene centrality, we generated ground truth gene centrality estimates as follows. We reasoned that a well-connected gene with high degree centrality would be highly co-expressed with many other genes in the genome, either in a linear or non-linear way. One way to quantitatively measure the degree of co-expression of a single query gene to all other genes is to measure how well the query gene can predict the expression level of all other genes in the genome. Therefore, our ground-truth gene centrality is defined as the percentage of variance explained by a query gene, with respect to all other genes in the genome. To measure percentage variance explained, for each gene in the genome, we trained a neural network consisting of a single input node (corresponding to the query gene expression), 3 hidden layers with 128, 512, and 1024 nodes, and a final output layer of 2000 nodes for all remaining genes in the genome. The percentage variance explained per gene was measured as 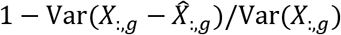 where 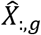 is a vector of expressions for *X*_:,*g*_ predicted by siVAE. We then averaged the percentage variance explained over all predicted genes, for all cells in a dataset, and refer to this quantity as the ground truth gene centrality.

### Identifying gene co-expression network neighbors from siVAE models

GCN inference methods typically output a weighted adjacency matrix that indicates the strength of co-expression of every pair of genes, which in turn could be used to find the closest neighbors of every gene in the genome based on the largest values of the adjacency matrix. In our experiments in **Figure 5c**, we used siVAE to also identify the closest co-expression neighbors of every gene in the genome, using two different approaches based on leveraging the feature embedding space. In our first approach (‘distance-based network neighbors’), we used Euclidean distance in the feature embedding space as a measure of distance between two genes in the network; the *k* nearest neighbors of a given query gene were defined as the *k* genes with shortest distance to the query gene. Our second approach, termed the ‘GCN-based network neighbors’, involved passing a matrix of feature embeddings to a GCN inference method (CLR) as input in place of the typical gene expression matrix input, in order to infer a classic gene co-expression network. From this gene co-expression network, we extracted the nearest neighbors of every gene according to the strategy described below for GCN inference methods.

### Identifying gene co-expression network neighbors using GCN inference methods

For GCN inference methods, we used the output adjacency matrix to identify the closest 20 neighborhood genes per target gene based on largest pairwise weights for each gene.

### Benchmarking gene neighborhoods

We computed the accuracy of both siVAE to the GCN inference methods in terms of their ability to identify neighbor genes that are co-expressed. To do so, we applied each method to identify the 20 closest neighbor genes to the query gene. We then defined a prediction task in which the 20 neighbor genes were used as input to a neural network to predict the expression of the target query gene. We used a fully connected neural network consisting of 3 hidden layers each with 16, 8, and 4 nodes, in addition to the input layer (with 20 nodes corresponding to the 20 closest neighbors), and the output layer consisting of a single node for the query gene. Accuracy was defined as the percentage variance explained with respect to the query gene. We compared siVAE to the GCN methods based on a set of 152 query genes, which were identified by taking the intersection of siVAE and each GCN inference method’s top 500 highest centrality genes, to ensure that the query genes were of high degree centrality (and therefore should have many neighbor genes).

### Quantifying overlap in gene neighborhoods between siVAE and other methods

We also used two different strategies to gauge the overlap in the gene neighborhoods predicted by siVAE and each GCN inference method, defined as the 20 closest genes to every query gene. For percentage overlap, we measured the percentage of genes that overlapped between any two sets of neighborhood genes. For mean correlation, we measured the Pearson correlations with twosided test of gene expression between every pair of genes between two neighborhood gene sets for the same query gene, then averaged the 20*20 = 400 correlation values together to compute mean correlation.

### Generating cell line embeddings with siVAE and graph kernel

We first divided the dataset by cell lines to generate a dataset per cell line. To avoid the downstream effect of datasets with different sizes, we performed downsampling by splitting each cell line into equal sized bins of 1000 cells. Bins that were not fully filled up were discarded for further analysis. Each binned datasets were fed into siVAE for generating gene embeddings and siVAE-inferred degree centrality. We then used in Weisfeiler-Lehman GraKeL^111^ to infer adjacency matrix per gene embedding then generate similarity matrix between inferred network generated per binned datasets. The downsampled versions of the same cell lines were averaged together in the similarity matrix before PCA visualization of the cell lines.

### Isolating and visualizing change in mitochondrial genes connectivity

Spearman correlation was measured between inferred degree centrality and differentiation efficiency per gene with Benjamini Hochberg correction for the P-values. For geneset enrichment analysis (GSEA) of the mitochondrial genes, we manually added MT gene set consisting of all the genes starting with “MT-” to the curated genesets of KEGG pathways (http://www.gsea-msigdb.org/gsea/msigdb/collections.isp). GSEA was performed using prerank function from GSEAPY package on the Spearman correlation coefficients^109^. Spearman correlation between mean expression of a gene and differentiation efficiency was measured through averaging the expression of a gene also with Benjamini Hochberg correction for P-values. Finally, we used Wilcoxon Rank Sum test to detect which gene-to-gene connections between mitochondrial genes led to significance difference in efficiencies of cell lines based on the inferred adjacency matrix. The mitochondrial gene network was visualized using Biocircos^112^ with all mitochondrial genes but only the significant mitochondrial edges.

### Testing for mitochondrial mutations causal for differentiation efficiency

Variant call files (VCF) per iPSC cell lines in NeuriDiff dataset were downloaded from HipSci data browser (https://www.hipsci.org/lines/#/lines). We filtered for only mitochondrial variants by discarding all variants not found in MT chromosome. We performed Wilcox rank sum test on individual variants with respect to differentiation efficiency by treating the cell lines either with or without variant as two independent groups. Next, we performed gene-based burden testing where the variants were grouped into genes based on genetic region and jointly correlated with neuronal differentiation efficiency. In addition to grouping based on individual mitochondrial genes, we also added additional grouping for all mitochondrial variants. Benjamini-Hochberg was used to correct for multiple testing for all tests.

## DECLARATIONS

siVAE has been uploaded to the PyPi software repository and can be found here: https://pypi.org/project/siVAE/.

The author(s) declare(s) that they have no competing interests.

